# benchmarkR: an R package for benchmarking genome-scale methods

**DOI:** 10.1101/018200

**Authors:** Xiaobei Zhou, Charity W. Law, Mark D. Robinson

## Abstract

**Summary:** **benchmarkR** is an R package designed to assess and visualize the performance of statistical methods for datasets that have an independent truth (e.g., simulations or datasets with large-scale validation), in particular for methods that claim to control false discovery rates (FDR). We augment some of the standard performance plots (e.g., receiver operating characteristic, or ROC, curves) with information about how well the methods are calibrated (i.e., whether they achieve their expected FDR control). For example, performance plots are extended with a point to highlight the power or FDR at a user-set threshold (e.g., at a method’s *estimated* 5% FDR). The package contains general containers to store simulation results (SimResults) and methods to create graphical summaries, such as receiver operating characteristic curves (rocX), false discovery plots (fdX) and power-to-achieved FDR plots (powerFDR); each plot is augmented with some form of calibration information. We find these plots to be an improved way to interpret relative performance of statistical methods for genomic datasets where many hypothesis tests are performed. The strategies, however, are general and will find applications in other domains.

**Availability:** The **benchmarkR** package is available from: https://github.com/markrobinsonuzh/benchmarkR

## 4 Introduction

The burden of proof in developing new statistical methods for inferring differences (e.g., changes in abundance) in genomic datasets is improved performance against existing methods. Methodologists typically resort to simulations since there is limited availability of large-scale validation datasets. To evaluate simulation performance (or performance with sufficient validation information), various metrics and plots are typically used, including but not limited to receiver operating characteristics (ROC) curves, which shows the tradeoff between true positive rates (TPR, or sensitivity, or power) and false positive rate across many cutoffs [1, 2], or false discovery (FD) plots, which highlight the cumulative number of false discoveries amongst the top ranked features.

While a method’s ability to give a good ranking is important, statistical methods typically build in some kind of adjustment to control the rate of errors made; in genomics, this typically takes the form of false discovery rate (FDR) control. Therefore, in these settings, it is of interest not only to know about detection performance (i.e., how well a statistical method separates true changes from false), but if and how well the error is controlled. To allow ourselves and the community more flexible ways to visualize additional information with respect to “calibration” in standard plots, such as ROC and FD plots, we developed the R-based **benchmarkR** package. In particular, we promote the use of a new variation: *power-to-achieved-FDR* plots at a small number of typical thresholds to directly contrast detection performance and error control. We find this plot to be the simplest way to digest both angles.

Figure 1 gives a simple but illustrative example. Suppose there is a simulation with a total of 10,000 features, of which 1,000 are truly differential. In this toy example, all features were generated for 3 replicates versus 3 replicates from a normal distribution with mean 0 and variance 1, except for the 1,000 differential features, which had a shifted mean (R code is available as Supplementary Material). If a method happens to systematically under- or overestimate the variance, this will lead to some initial clues in the distribution of raw P-values, but it may not compromise the method’s ability to rank differential features. A *calibrated* method should show a mixture of uniform P-values with the differential features showing strong statistical evidence as a peak at the low end (Figure 1a). A (systematically) *liberal* method will tend to overstate the statistical evidence and push all P-values toward 0 (Figure 1b), whereas a *conservative* method will push P-values towards 1 (Figure 1c), relative to a calibrated method (in our toy example, this calibration is modified through the variance estimates). In practice, P-value distributions may be hard to diagnose since a combination of factors will affect their overall shape and other problems may arise, such as correlation of observations, model misspecification or outliers. Importantly, calibration is not well represented in an ROC curve (Figure 1d). Despite the differences in statistical calibration that we have introduced to the toy example, the ROC curve cannot relay any difference in performance (in the toy example, the calibration does not strongly affect the ranking). In addition, ROC curves can actually be misleading because it is not known where a particular usage of a method (e.g., FDR=5%) will lie on the curve. For example, a method could have a great ability to rank features (a high ROC curve and area under the curve), but it may be extremely conservative and thus not very useful in practice. The ROC curve plotted using the **benchmarkR** package (Figure 1d; rocX method) highlights the point on each ROC curve that corresponds to the method’s estimated 5% FDR threshold; however, it is important to note that the method does not necessarily achieve this level of control. An alternative method to look at simulation results is an FD plot (Figure 1e), where the cumulative number of false discoveries is displayed amongst the top ranked differential features; fewer FDs are desirable. The **benchmarkR** package provides a variation of this plot (using the fdX method) that adds the location of the method’s operating position (e.g., FDR=5%). This allows the methodologist to get a sense of whether methods are adequately controlling their FDR. Pushing this further, we find a power-to-achieved-FDR plot (via the powerFDR method) to be a concise summary of both angles (Figure 1f). In this plot, several typical FDR thresholds are used (e.g., FDR=1%, 5%, 10%) and for each threshold, the method’s performance in terms of power and achieved FDR are plotted, with a line joining the different cutoffs. For these plots, it is desirable when the method is able to control the FDR, which would require the method’s X-axis point to be on the left side of the corresponding threshold line; if this occurs, the default plotting system in **benchmarkR** will use a filled-in symbol whereas if the error is not controlled, an open symbol will be used.

**Figure 1:**
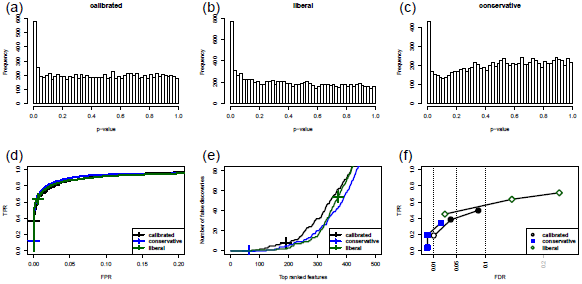

A hypothetical example of a calibrated, liberal and conservative statistical method in genomics. Panels (a)-(c) should P-value distributions. Panels (d), (e), (f) show an ROC curve (`rocX`), a false discovery plot (`fdX`) and a power-versus-achieved-FDR plot (`powerFDR`), respectively. The code to regenerate this plot is available as Supplementary Material.

## 5 Implementation

A typical use of the **benchmarkR** package may look like the following:

~~~
library("benchmarkR")

# create container for results re <- SimResults(pval,labels)
# 3-panel plot

benchmarkR(re)

# individual plots
rocX(re)
fdX(re)
powerFDR(re)
~~~

where `pval` (a vector or matrix) and `labels` (a vector) give the scores and labels, respectively. The `benchmarkR` function is simply a wrapper that makes a 3-panel plot consisting of `rocX`, `fdX` and `powerFDR`. Each individual plot is highly customizable; see the package vignette for further details.

## Acknowledgement

The authors wish to thank all members of the Robinson laboratory for helpful discussions. We wish to acknowledge funding from an SNSF Project Grant (143883) and the European Commission through the 7th Framework Collaborative Project RADIANT (Grant Agreement Number: 305626).

Conflict of Interest: none declared.

